# Unlocking the Mechanism of Action: A Cost-Effective Flow Cytometry Approach for Accelerating Antimicrobial Drug Development

**DOI:** 10.1101/2023.11.07.566019

**Authors:** Fabian Mermans, Hanna De Baets, Cristina García-Timermans, Wim Teughels, Nico Boon

## Abstract

Antimicrobial resistance is one of the greatest challenges to global health. While the development of new antimicrobials can combat resistance, low profitability is reducing the number of new compounds brought to the market. Elucidating the mechanism of action is crucial for developing new antimicrobials. This process can become expensive as there are no universally applicable pipelines and scientific expertise in different fields is required. One way to determine the mechanism of action is the use of predictive modeling, as antimicrobials can be classified into limited groups.. We demonstrate a cost-effective flow cytometry approach for determining the mechanisms of action of new compounds. Cultures of *Actinomyces viscosus* and *Fusobacterium nucleatum* were treated with a range of antimicrobials and measured by flow cytometry. A Gaussian mixture mask was applied over the data to construct a phenotypic fingerprint. The fingerprints were used to train random forest classifiers, and classifiers were used to predict the mechanism of action of cephalothin. Significant statistical differences were found among the 10 different treatment groups. A pairwise comparison between treatment groups showed a statistical difference for 35 out of 45 pairs for *Actinomyces viscosus* and 32 out of 45 pairs for *Fusobacterium nucleatum* after 3.5h of treatment. The best performing random forest classifier yielded a Matthews correlation coefficient of 0.92 and the mechanism of action of cephalothin could be successfully predicted. These findings suggest that flow cytometry can be a cheap and fast alternative for determining the mechanism of action of new antimicrobials.

## Introduction

The use of antibiotics and antiseptics has revolutionized modern medicine over the past century by allowing us to treat previously lethal diseases (CDC 1999; Armstrong, Conn, and Pinner 1999). However, emerging antimicrobial resistance (AMR) is currently one of the greatest challenges in global health. For 2019, it is estimated that 4.95 million deaths were associated with bacterial AMR globally (Murray et al. 2022). AMR is a consequence of natural evolution as well as the improper use of antimicrobials (Bell et al. 2014; Lipsitch and Samore 2002). One of the few ways to tackle AMR is through the development of new antimicrobials. Nonetheless, the number of new compounds brought to the market is declining (Cook and Wright 2022). Development is expensive, usually in the range of hundreds of millions of US dollars. At the same time, profitability is low, leading to a limited economic incentive to invest in new compounds (Boyd, Teng, and Frei 2021). One important step in the development of new compounds is the elucidation of the mechanism of action (MOA). Both the European Medicines Agency (EMA) and the U.S. Food and Drug Administration (FDA) state that, if possible, the MOA of a novel antimicrobial should be known (FDA 1985; European Parliament and Council of the European Union 2001). Even though it is not strictly necessary to elaborate the MOA to obtain approval, it can help in selecting which compounds to investigate further. When insight in the MOA is lacking, clinical trials are more likely to fail, resulting in higher costs. Additionally, the information can be used to modify molecules of interest to enhance their pharmacokinetic and pharmacodynamic properties (*e.g.* lower host toxicity) (Hudson and Lockless 2022; Drews 2000; Gregori-Puigjané et al. 2012). Usually, elucidation of the MOA is part of the pre-clinical development after hit-to-lead compound selection. Costs to determine the MOA can become significant as there are no universally applicable pipelines to do so and expertise in microbiology, genetics, chemical biology, genomics, and biophysics is required (Miethke et al. 2021). For example, techniques often used for molecular target identification are transcriptomics-based pattern recognition, metabolomics, FTIR spectroscopy, and specialized proteomics (O’Rourke et al. 2020; Hoerr et al. 2016; Altharawi, Rahman, and Chan 2019; Schopper et al. 2017).

In general, the MOA of antimicrobials can be classified into a limited number of groups. The first group targets the cytoplasmic membrane and aims to disrupt it. Examples include chlorhexidine and polymyxins. The second group tampers with cell wall synthesis, such as the β-lactams and glycopeptides. Both compounds belonging to the first and second groups are considered bactericidal. The third group is composed of compounds that interfere with nucleic acid synthesis. Here, it is possible to distinguish different subgroups: compounds that target replication (*e.g.* metronidazole) or compounds that target transcription (*e.g.* rifampicin). The fourth group are inhibitors of protein synthesis. Again, subgroups are distinguished: the compounds that target the 50S subunit of the bacterial 70S ribosome (*e.g.* macrolides, clindamycin, streptogramins), the ones targeting the 30S subunit of the bacterial 70S ribosome (*e.g.* tetracycline, aminoglycosides), and the ones that cause premature termination of translation (*e.g.* puromycin). The last group consists of compounds that inhibit folic acid metabolism (*e.g.* sulfonamides, trimethoprim) (Kapoor, Saigal, and Elongavan 2017; Aviner 2020; Lim and Kam 2008; Dingsdag and Hunter 2018).

The occurrence of a limited number of classes hints at the use of predictive modelling based on the MOA of proven compounds to assess the MOA of new compounds (Hoerr et al. 2016; Athamneh et al. 2014; Mongia, Guler, and Mohimani 2022). One way to do so is to assess the phenotypic state of microbial cells after exposure to different antimicrobials. Flow cytometry is one such technique in which the phenotypic attributes of microbial cells can be assessed in a high throughput manner (Müller and Nebe-von-Caron 2010; Hatzenpichler et al. 2020). Additionally, the measurement of a single sample is cheap, usually costing less than 1 euro. The information obtained from flow cytometric measurements can be used to build a phenotypic fingerprint of the microbial population and is suitable for the study of phenotypic heterogeneity between samples (Props et al. 2016; Koch et al. 2013; De Roy et al. 2012). The discriminative power of phenotypic fingerprints can be increased using adaptive binning approaches, such as PhenoGMM (Rubbens et al. 2021a).

In this study, we explored the use of flow cytometric microbial fingerprinting to predict the MOA of antimicrobial compounds. We exposed axenic cultures of *Fusobacterium nucleatum* and *Actinomyces viscosus* to a wide range of antimicrobials and performed flow cytometric measurements. The flow cytometry data were used to construct phenotypic fingerprints based on Gaussian mixture models. In turn, these fingerprints were used to train random forest classifiers. Furthermore, we investigated if clustering of MOA classes could be observed after treating a complex microbial community (*i.e.* saliva) with different antimicrobials. Finally, trained classifiers were used to predict the MOA of an unseen compound, cephalothin.

## Materials and methods

### Strains and growth conditions

*Actinomyces viscosus* ATCC 15987 and *Fusobacterium nucleatum* ATCC 10953 were maintained on blood agar base No. 2 (Oxoid, Hampshire, UK) supplemented with menadione (1 mg/mL) (Sigma Aldrich, Diegem, Belgium), hemin (5 mg/mL) (Sigma Aldrich, Diegem, Belgium), and 5% sterile defibrinated horse blood (Oxoid, Hampshire, UK) under anaerobic conditions (90% N_2_, 10% CO_2_) at 37°C. Both strains were incubated for 16 h before treatment in Brain Heart Infusion (BHI) broth (Karl Roth, Karlsruhe, Germany) under anaerobic conditions (90% N_2_, 10% CO_2_) at 37°C.

### Treatment

After 16h of growth, bacterial cultures were divided into aliquots and supplemented with one antimicrobial compound in triplicate (final volume: 300 µL). An untreated sample was included as a control in triplicate. Consecutively, samples were incubated under anaerobic conditions (90% N_2_, 10% CO_2_) at 37°C for 3.5 h or 24 h. Table 1 shows tested antimicrobial compounds with the respective concentrations in which they were supplemented with the bacterial samples.

**Table 1.**
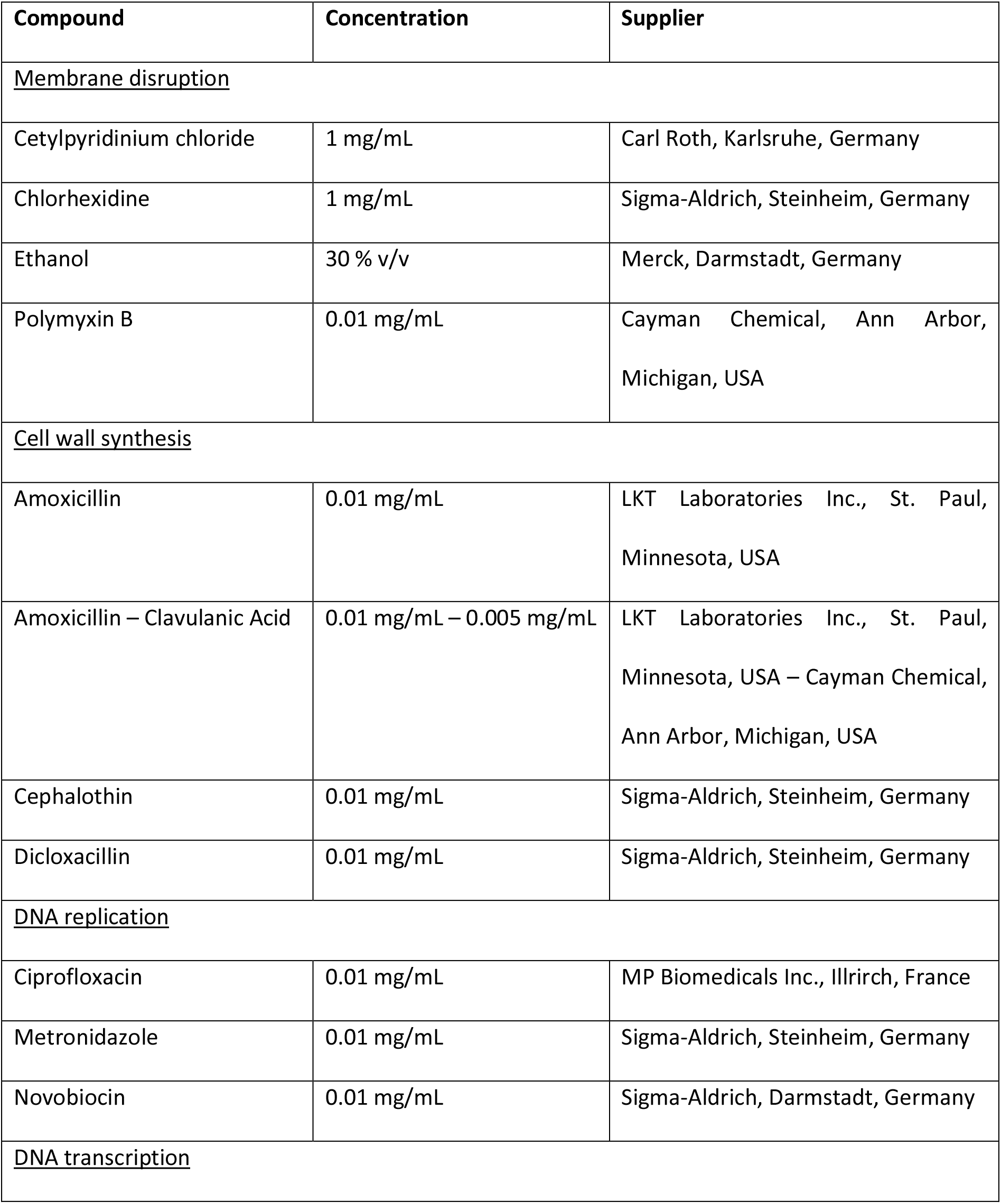
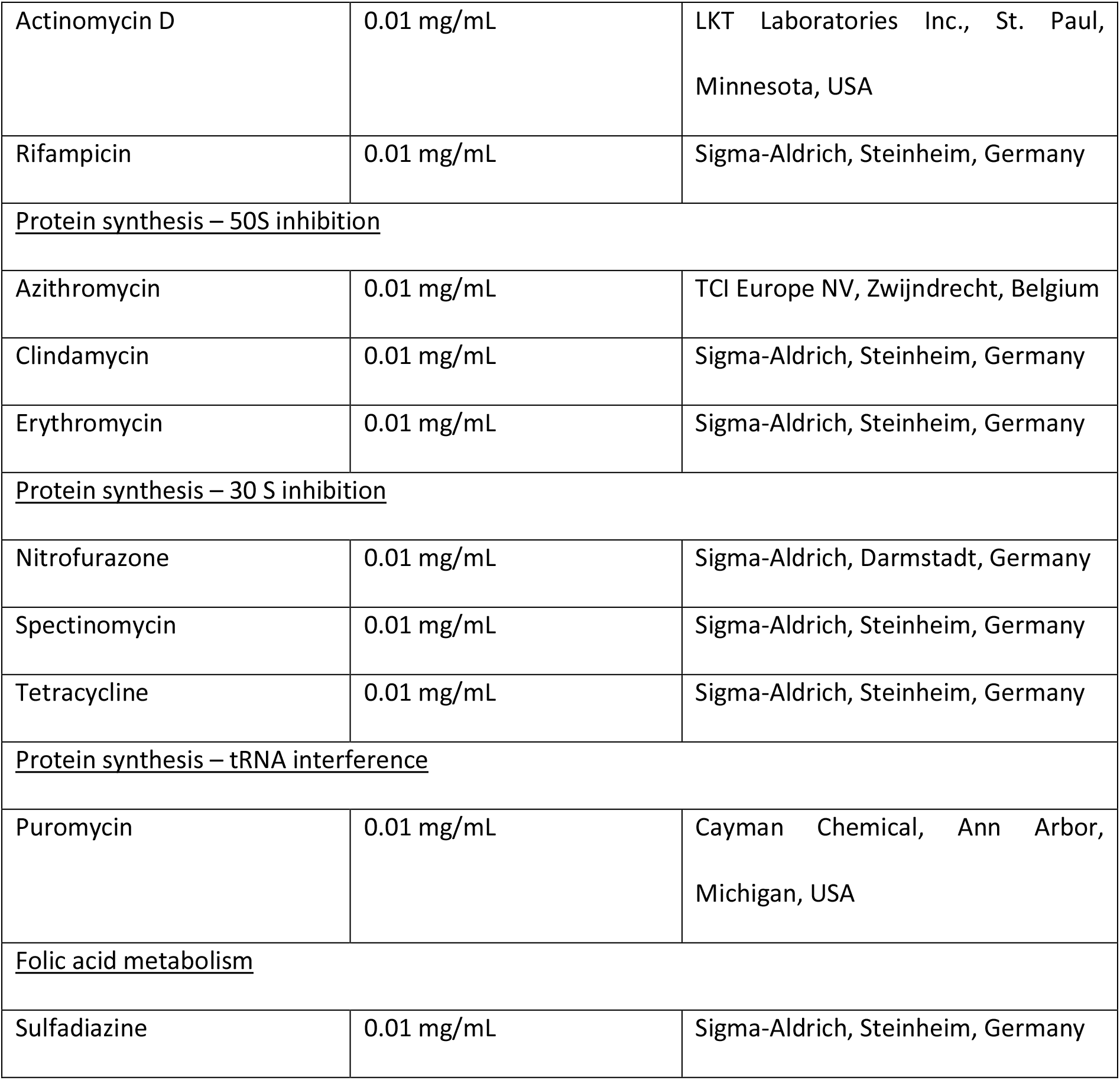
Antimicrobial compounds with their respective final concentration and supplier, grouped by mechanism of action.

After incubation with the antimicrobials, samples were diluted 100x in sterile and 0.2 µm filtered phosphate buffered saline (PBS) (PBS tablet, Sigma-Aldrich, Steinheim, Germany) in order to stop the treatment. Additionally, both an untreated sample and a heat-treated (40 min at 110°C) sample without consecutive incubation were included as controls in triplicate.

### Saliva sampling and treatment

Saliva was collected from a healthy individual (male, 28 years) by passive drooling for 5 minutes in a sterile container. The donor was asked to refrain from brushing teeth, using mouth rinses, and eating and drinking for two hours prior to the donation. Consecutively, saliva was diluted 10x in sterile and 0.2 µm filtered phosphate buffered saline (PBS) (PBS tablet, Sigma-Aldrich, Steinheim, Germany). The diluted sample was then divided into aliquots that were each supplemented with one antimicrobial compound in triplicate (final volume: 300 µL). The supplemented antimicrobials were amoxicillin, amoxicillin-clavulanic acid, azithromycin, ciprofloxacin, clindamycin, erythromycin, and metronidazole (Table 1). Non-supplemented 10x diluted saliva served as a control and was included in triplicate. Next, samples were incubated under aerobic conditions at 37°C for 24h. After incubation, samples were diluted 100x in sterile and 0.2 µm filtered PBS to stop the treatment. The salivary donor signed an informed consent before the donation. The study was approved by the Ethics Committee of the University of Ghent (B6702022000406).

### Flow cytometric measurements

#### Sample preparation

Directly after stopping the treatment, samples were stained with SYBR® Green I/propidium iodide (SGPI) and analyzed using flow cytometry in order to determine the phenotypic fingerprint and assess membrane integrity(Grégori et al. 2001). SYBR® Green I (10 000x concentrate in DMSO) (Invitrogen, Eugene, USA) and propidium iodide (20mM) (Invitrogen, Eugene, USA) were diluted 100 and 50 times, respectively, in 0.2 µm filtered DMSO (Merck, Darmstadt, Germany). Samples were then stained with 1% v/v SGPI and incubated in the dark at 37°C for 20 minutes.

#### Sample measurement

Stained samples were measured using an Attune NxT (Invitrogen, Carlsbad, USA) flow cytometer equipped with a blue (488 nm) and red (638 nm) laser. Performance of the instrument was checked using Attune Performance tracking beads (Invitrogen, Eugene, USA). Only the blue laser was used for the excitation of the stains. A 530/30 nm band-pass filter was used for the detection of green fluorescence (BL1) and a 695/40 nm band-pass filter was used for the detection of red fluorescence (BL3). A 488/10 band-pass filter was used for the detection of forward scatter (FSC) and side scatter (SSC). The flow rate was set to 100 µL/min and stop conditions were set to 100 µL of sample analyzed. The threshold (set on green fluorescence, BL1) and PMT-voltages were determined by using control samples: an untreated sample, a heat-killed sample (110 °C for 40 minutes), sterile BHI, and sterile PBS.

#### Data analysis

Flow cytometry data were imported in R (version 4.2.0) using the flowCore package (version 2.8.0) (Hahne et al. 2009). Data were transformed using the arcsine hyperbolic function (Props et al. 2016), and gated manually on the primary fluorescent channels (BL1 and BL3) to remove background (Supplementary 1). Consecutively, data were normalized by dividing each parameter by the maximum observed value for SYBR® Green I fluorescence (BL1) over all treatments within one bacterial strain and treatment duration, for each bacterial strain and for each treatment duration. The same was done for the saliva samples.

Application of a Gaussian mixture mask to identify clusters within the flow cytometry data was performed using the ‘*PhenoGMM*’ function of the Phenoflow package (version 1.1.2) (Rubbens et al. 2021a). To be able to generate the mask, samples were pooled based on their MOA class and consecutively subsampled to an equal number of cells per class. This was done to avoid biased model training towards a specific class. The Gaussian mixture model (GMM) was optimized using the Bayesian information criterion (BIC) (Ludwig et al. 2019; Heyse et al. 2021). The GMM leads to a one-dimensional (1D) vector for each sample that represents the number of cells allocated to each cluster in the model. The parameters upon which the model was build were FSC-H, FSC-A, SSC-H, SSC-A, BL1-H, BL1-A, BL3-H, and BL3-A because these are expected to contain the most information (Rubbens et al. 2017). Model output was used to perform both NMDS (vegan package version 2.6-2) (Oksanen et al. 2022) and PCoA (vegan package version 2.6-2) (Oksanen et al. 2022) in order show robustness in observations. Hierarchical cluster analysis was done using the ‘*hclust*’ function of the stats package (version 4.2.2). For the axenic bacterial cultures, statistical differences between mechanism of action classes were determined through distribution independent analysis of similarities (ANOSIM) (vegan package version 2.6-2) (Oksanen et al. 2022). P-values for pairwise comparisons were adjusted using the Benjamini and Hochberg method (stats package version 4.2.2).

Random forest classifiers to predict the mechanism of action were trained on PhenoGMM output of the axenic bacterial strains using the caret package (version 6.0-92) (Kuhn et al. 2022). The Matthews correlation coefficient (MCC) was used as a performance metric to optimize the classifiers. The MCC was used to account for class imbalance and was calculated using the mltools package (version 0.3.5) (Gorman 2018). The number of trees was set to 500 for all models. All models were trained using a 3-times repeated 5-fold cross validation scheme. The classification performance of the classifiers was assessed using a 5-fold nested 3-times repeated 5-fold cross validation scheme. Ultimately, classifiers were used to predict the mechanism of action of a compound that was not used in the training and testing data (*i.e.* cephalothin).

## Results

### Diversity of antimicrobial treated samples

Axenic cultures of *A. viscosus* and *F. nucleatum* were treated with antimicrobials for 3.5h or 24h and consecutively measured by flow cytometry. Membrane integrity was assessed through manual gating (Supplementary 1). An increase in cell population with damaged membranes is observed with increased treatment time for most antimicrobials for *A. viscosus* and for some antimicrobials for *F. nucleatum* (Supplementary 2). Phenotypic fingerprints were constructed by applying a Gaussian mixture mask to the flow cytometric data (Supplementary 3). Grouping according to MOA class can be observed based on the phenotypic fingerprint using NMDS for *F. nucleatum* and *A. viscosus* after 3.5h and 24h of treatment (Figure 1). Clear separate groups for MOA classes ‘Membrane Disruption’ and ‘Cell Wall Synthesis’ are visible for *A. viscosus*, except for polymyxin B, which is separate from the ‘Membrane Disruption’ class. For *F. nucleatum,* distinctive groups were only observed for the ‘Membrane Disruption’ class at both treatment durations and for the ‘Cell Wall Synthesis’ class after 24h of treatment. Robustness in observations was demonstrated by PCoA for both bacterial strains (Supplementary 4). Grouping was driven by strain when both strains were considered together, especially for the shorter treatment duration (Supplementary 5). For *A.* viscosus, cluster analysis revealed separate clusters for the classes ‘Membrane Disruption’, ‘Cell Wall Synthesis’ and ‘Protein Synthesis: 50S Inhibition’ after 3.5h of treatment. After 24h of treatment, only the class ‘Cell Wall Synthesis’ formed a separate cluster. As with the ordination, polymyxin B was not clustering with its respective class at both treatment times. For *F. nucleatum*, only a separate cluster for the class ‘Cell Wall Synthesis’ could be observed after 24h of treatment (Figure 2).

**Figure 1.**
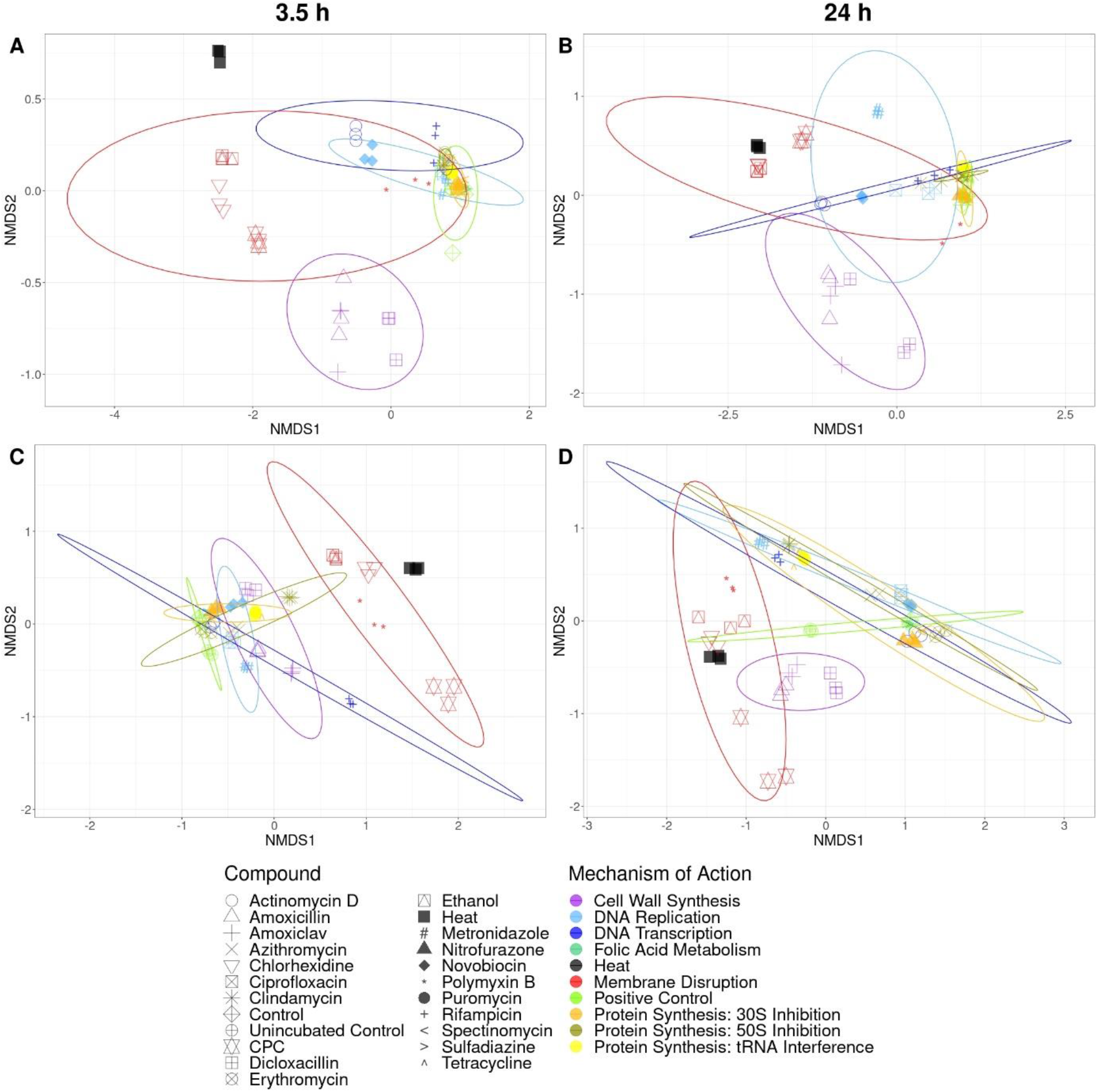
NMDS of flow cytometric fingerprints of *A. viscosus* (A, B) and *F. nucleatum* (C, D) after 3.5h (A, C) and 24h (B, D) of treatment with antimicrobials. Phenotypic fingerprints were generated using PhenoGMM (Phenoflow package). Compounds with the same MOA are grouped by color. Ellipses were drawn at the 95% confidence level. ‘Heat’ indicates the heat-treated control; ‘Control’ indicates the untreated sample that underwent incubation with the antimicrobial treated samples; ‘Unincubated Control’ indicates the untreated sample that did not undergo consecutive incubation.

**Figure 2.**
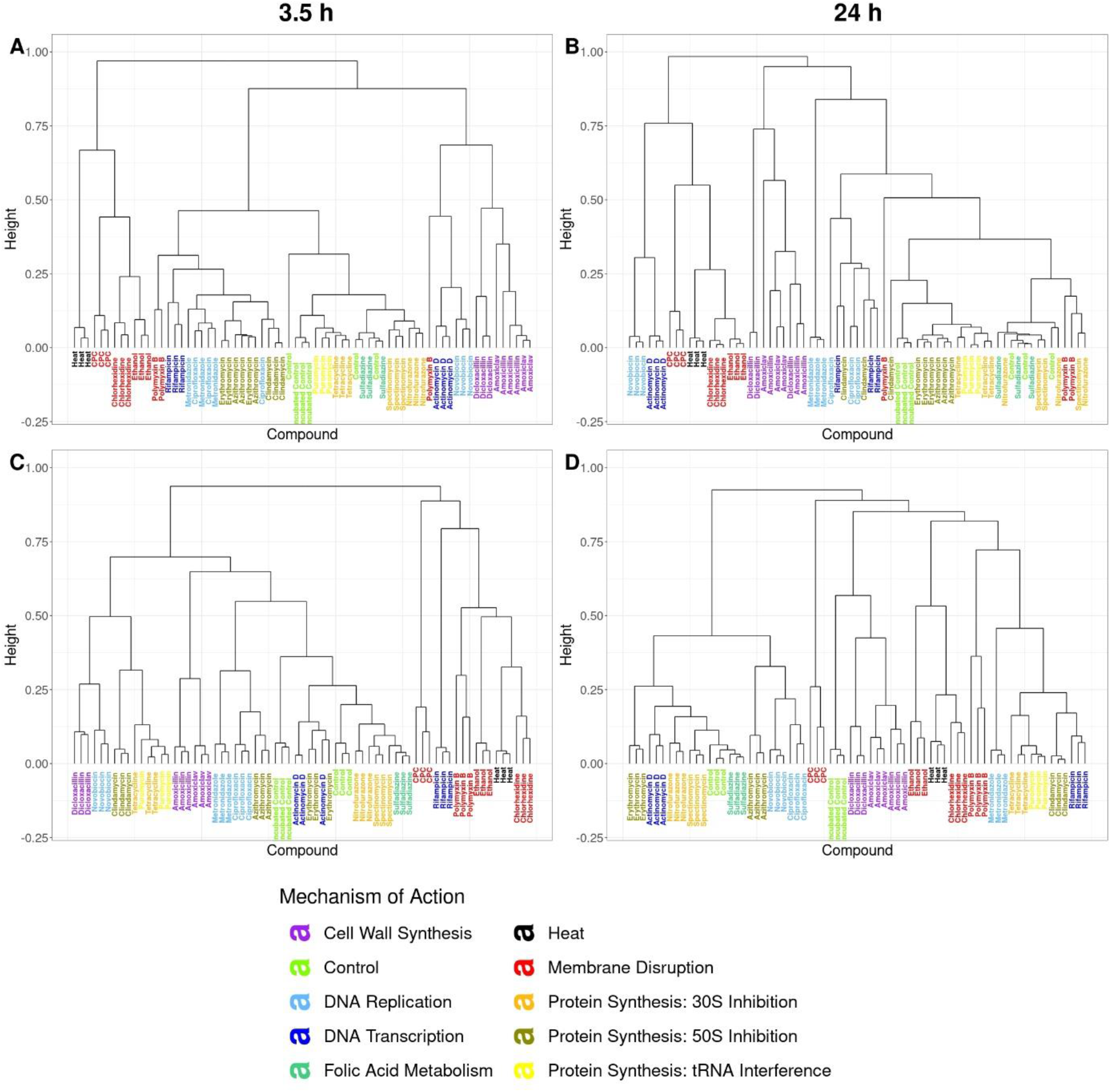
Dendrograms of flow cytometric fingerprints of *A. viscosus* (A, B) and *F. nucleatum* (C, D) after 3.5h (A, C) and 24h (B, D) of treatment with antimicrobials. Compounds with the same MOA are grouped by color.

Significant statistical differences between phenotypic fingerprints of different MOA classes were found for both *A. viscosus* (R = 0. 5625, p = 0.0001 at 3.5h; R = 0.5135, p = 0.0001 at 24h) and *F. nucleatum* (R = 0.4764, p = 0.0001 at 3.5h; R = 0.3763, p = 0.0001 at 24h). For *A. viscosus*, pairwise comparison between MOA classes showed a significant statistical difference for 35 out of 45 pairs after 3.5h of treatment with antimicrobials and for 34 out of 45 pairs after 24h of treatment (p < 0.05; Supplementary 6). For *F. nucleatum*, 32 out of 45 pairs after 3.5h of treatment and 18 out of 45 pairs after 24h of treatment were significantly different (p < 0.05; Supplementary 6). The ‘Cell Wall Synthesis’ class was significantly different from all other classes for both strains for both treatment durations. ‘Membrane Disruption’ was different from all other classes except for ‘Heat’ for both strains at both treatment durations, for ‘DNA Transcription’ for *A. viscosus* after 24h of treatment and for ‘Protein Synthesis – tRNA Interference’ for *F. nucleatum* after 24h of treatment. ‘DNA Replication’ and ‘DNA Transcription’ classes were different from most other classes for both strains after 3.5h of treatment and for *A. viscosus* after 24h of treatment. For *F. nucleatum,* the different ‘Protein Synthesis’ classes could not be distinguished from each other. However, for *A. viscosus* ‘30S Inhibition’ and ‘50S Inhibition’, as well as ‘50S Inhibition’ and ‘tRNA Interference’ were significantly different after 3.5h of treatment. ‘50S Inhibition’ and ‘tRNA Interference’ were not significantly different from each other anymore after 24h of treatment. The ‘Folic Acid Metabolism’ and ‘Protein Synthesis – tRNA Interference’ classes showed the least significant differences from other classes. Moreover, they did not differ from the control, except for *F. nucleatum* after 3.5h of treatment and for the ‘Protein Synthesis – tRNA Interference’ class for *A. viscosus* after 24h of treatment.

A saliva sample was treated with different antimicrobials for 24h and measured using flow cytometry. Phenotypic fingerprints were constructed by applying a Gaussian mixture mask to the flow cytometric data (Supplementary 3). NMDS reveals grouping according to MOA class based on the phenotypic fingerprint of saliva (Figure 3). Grouping is most prominent for the ‘Cell Wall Synthesis’ class and the ‘DNA Replication’ class. Cluster analysis shows no distinct clusters between MOA classes (Figure 3).

**Figure 3.**
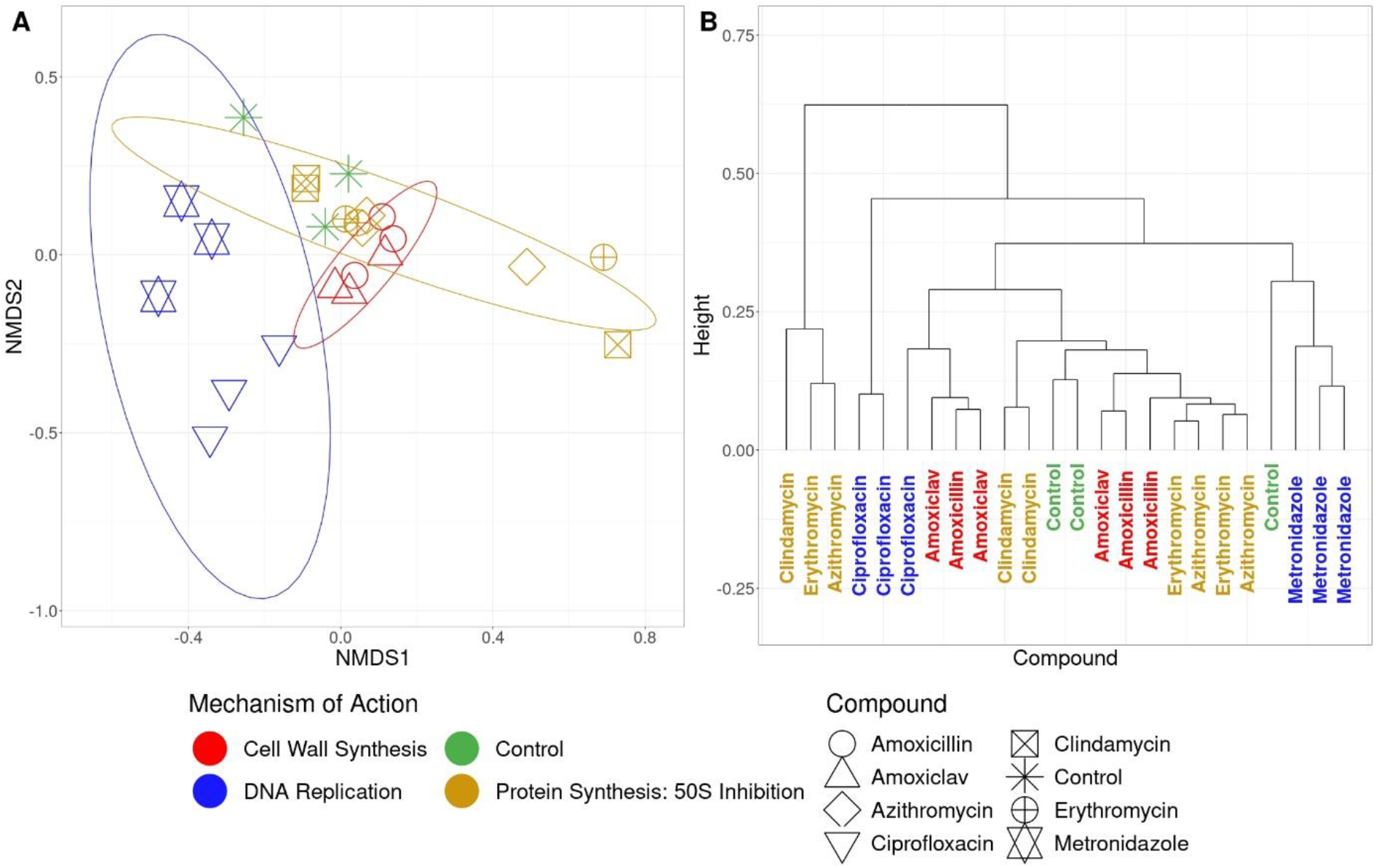
Diversity analysis of phenotypic fingerprints of a saliva sample treated with different antimicrobials for 24h. Compounds with the same MOA are grouped by color. ‘Control’ indicates to the untreated sample that underwent incubation with the antimicrobial treated samples. (A) NMDS of flow cytometric fingerprints. Ellipses were drawn at the 95% confidence level. (B) Dendrogram of flow cytometric fingerprints.

### Random Forest Classification

Random forest classifiers could be successfully trained using the PhenoGMM-generated fingerprints for *A. viscosus* and *F. nucleatum* at both treatment durations (Table 2). Classifiers trained using *F. nucleatum* as a model strain performed better compared to classifiers trained with *A. viscosus*. The influence of the duration of treatment was inversed between bacterial strains.

**Table 2.**
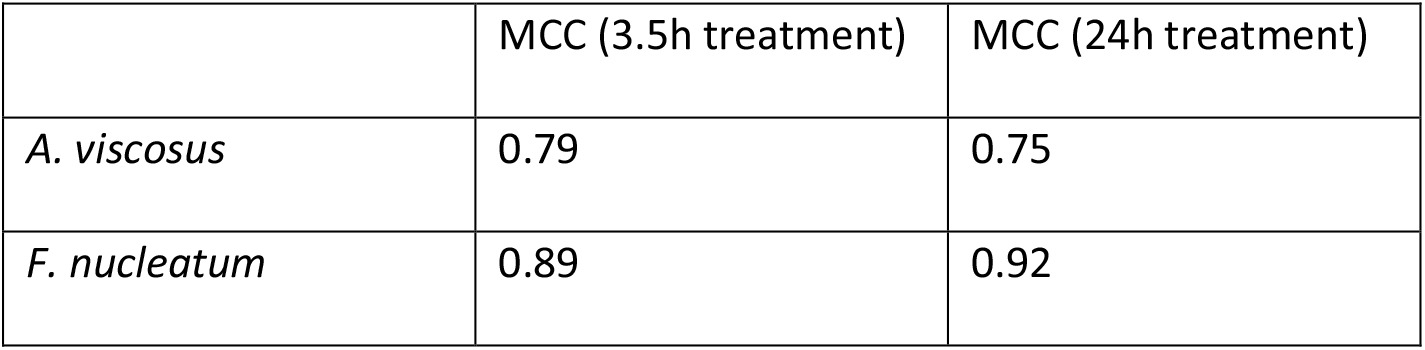
Matthews correlation coefficients (MCC) of trained random forest classifiers for *A. viscosus* and *F. nucleatum* after 3.5h and 24h of treatment with the antimicrobials. Classifiers were trained using the phenotypic fingerprints generated with PhenoGMM. The MCC was used as performance metric to optimize the model to account for class imbalance. A MCC of ‘0’ indicates random guessing; a MCC of ‘1’ indicates perfect classification.

MOA classes ‘Membrane Disruption’, ‘Cell Wall Synthesis’ and ‘DNA Replication’ can be predicted with high accuracy (>87.5%) for both bacterial strains (Table 3). Additionally, high accuracy (> 92.3%) was observed for the ‘Protein Synthesis – 50S Inhibition’ class, except for *F. nucleatum* after 3.5h of treatment.

**Table 3.** Performance of each MOA class for the different random forest classifiers trained using the phenotypic fingerprints of antimicrobial-treated cultures of *A. viscosus* and *F. nucleatum*.

|  | *A. viscosus* |  |  | *F. nucleatum* |  |
| --- | --- | --- | --- | --- | --- |
|  | 3.5h<br>treatment | 24h<br>treatment |  | 3.5h<br>treatment | 24h<br>treatment |
| Class | Accuracy<br>(%) | Accuracy<br>(%) |  | Accuracy<br>(%) | Accuracy<br>(%) |
| Protein Synthesis tRNA Interference | 25 | 75 |  | 100 | 100 |
| Protein Synthesis 50S Inhibition | 100 | 92.3 |  | 69.2 | 100 |
| Protein Synthesis 30S Inhibition | 92.3 | 69.2 |  | 92.3 | 76.9 |
| Control | 66.7 | 75 |  | 100 | 100 |
| Membrane Disruption | 87.5 | 87.5 |  | 100 | 100 |
| Heat | 100 | 100 |  | 75 | 75 |
| Folic Acid Metabolism | 25 | 0 |  | 100 | 100 |
| DNA Transcription | 50 | 75 |  | 87.5 | 75 |
| DNA Replication | 92.3 | 92.3 |  | 92.3 | 100 |
| Cell Wall Synthesis | 100 | 100 |  | 100 | 100 |

Confusion matrices reveal that incorrect predictions are not distributed evenly over all classes (Supplementary 7). For *A. viscosus* after 3.5h of treatment, the ‘DNA Transcription’ class was most often confused with the ‘Protein Synthesis – 50S Inhibition’ class, the ‘Folic Acid Metabolism’ class was confused most often with the ‘Control’ class, and vice versa, and the ‘Protein Synthesis – tRNA Interference’ class was confused most often with the ‘Protein Synthesis – 30S Inhibition’ class. After 24h of treatment, the ‘Protein Synthesis – 30S Inhibition’ class was mostly confused for being the ‘Protein Synthesis – 50S Inhibition’ class, and the ‘Folic Acid Metabolism’ class was most often confused with the ‘Protein Synthesis – 30S Inhibition’ class. For *F. nucleatum,* the ‘Protein Synthesis – 50S Inhibition’ class was mostly misclassified as the ‘DNA Transcription’ class after 3.5h of treatment, and the ‘Protein Synthesis – 30S Inhibition’ class was mostly misclassified as the ‘Protein Synthesis – tRNA Interference’ class after 24h of treatment.

### Prediction of MOA of an unseen compound

The MOA of the unseen compound cephalothin could be successfully predicted as the ‘Cell Wall Synthesis’ class by most classifiers (Table 4). Only for the classifier trained on *F. nucleatum* after 24h of treatment, two of the replicates were predicted incorrectly as the ‘Membrane Disruption’ class. However, for these replicates, the second most likely class was ‘Cell Wall Synthesis’ with a probability of 38.6% (replicate 2) and 38.6% (replicate 3).

**Table 4.**
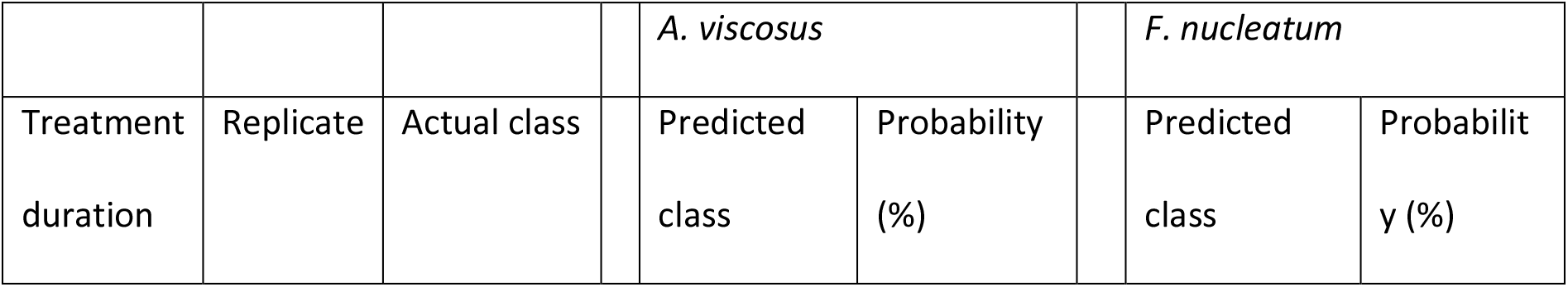
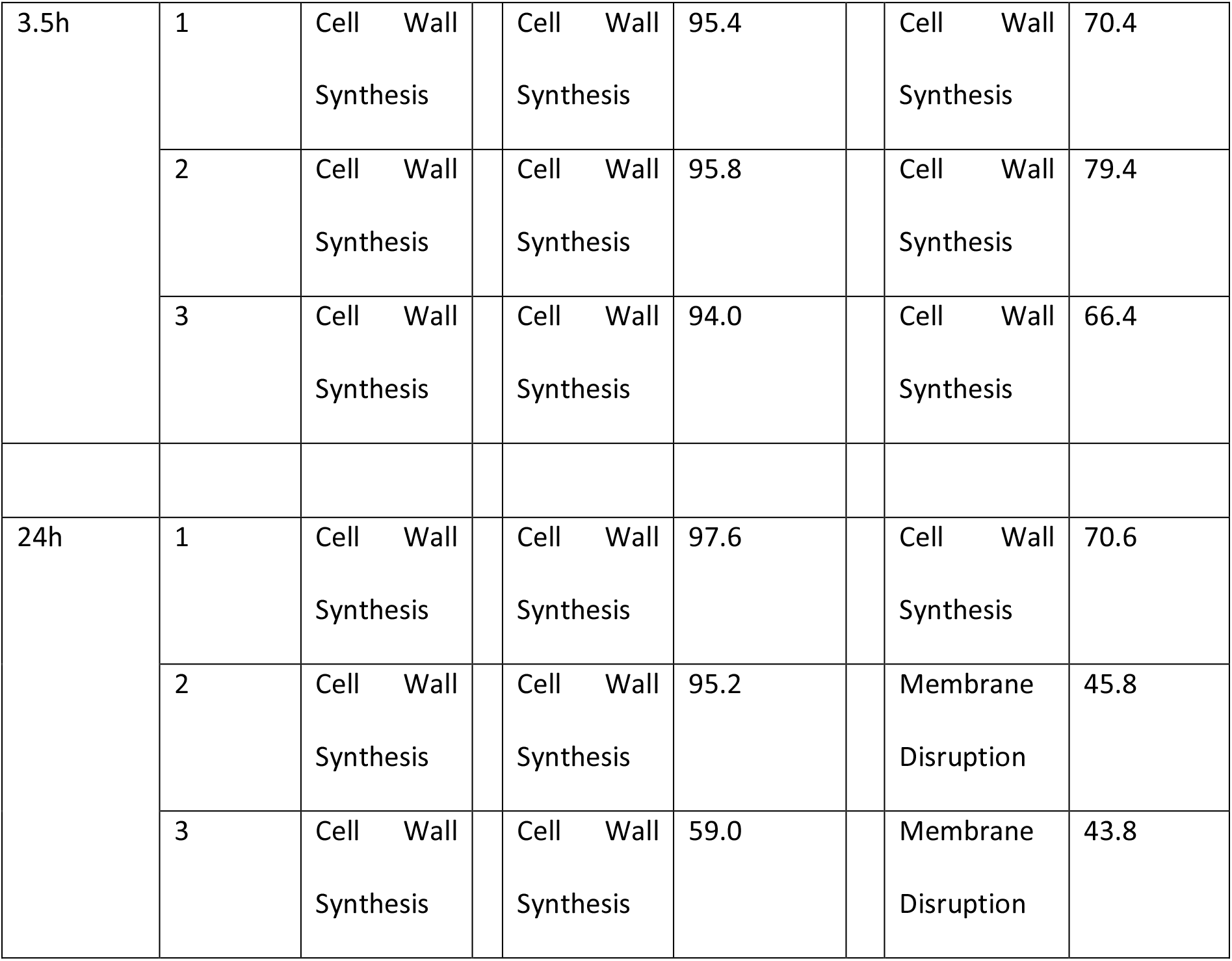
Predicted MOA classes for the unseen compound cephalothin for random forest models based on the PhenoGMM fingerprint. Predictions were made for each individual replicate.

## Discussion

The number of novel antimicrobials brought to market is constrained by high development costs and low profitability. The elucidation of the MOA can contribute significantly to this cost (Miethke et al. 2021). To overcome this obstacle, we investigated if flow cytometric microbial fingerprinting could be used as a fast and cheap alternative to predict the MOA of antimicrobials. Our experimental results show that the phenotypic fingerprints of different MOA groups could be distinguished from each other. Moreover, random forest classifiers predicting the MOA could be successfully trained using these fingerprints. Finally, trained random forest classifiers were able to predict the MOA of an unseen compound.

Phenotypic fingerprints of samples treated with antimicrobials cluster according to MOA class, with clustering of MOA classes that target the integrity of the cell being most prominent. This is the result of the use of SGPI to stain bacterial cells, as SGPI is used to assess membrane integrity (Grégori et al. 2001). Furthermore, the MOA class ‘DNA Replication’ that influences the level of DNA is shown to be statistically different from other MOA classes as well. Again, this may be the result of the staining since SYBR® Green I (SG) binds to double stranded DNA (Dragan et al. 2012). Therefore, the use of different stains may increase discriminative power between MOA classes. For example, the use of bio-orthogonal non-canonical amino acid tagging (BONCAT) could help in discriminating between the different ‘Protein Synthesis’ classes (Lindivat et al. 2020). Additionally, increased discriminative power could be achieved by combining multiple phenotypic fingerprints acquired with different staining techniques.

A lower number of statistically significant differences between MOA classes were found with increased treatment time for both bacterial strains (Supplementary 6). For *F. nucleatum,* it has to be noted that seven pairwise comparisons that were significantly different after 3.5h of treatment but not significant after 24h, were comparisons involving the untreated control. Figure 1 (D) shows a large difference between the untreated control that did not undergo incubation and the control that did, leading to high variability in this MOA group. This high within variability could explain the reduced statistical differences for these comparisons. For all other cases where statistical differences were not significant after 24h anymore, we argue that it is the result of phenotypes of bacterial cells becoming more similar with increased treatment time and that these phenotypes are related to cellular death. For example, for both *A. viscosus* and *F. nucleatum*, MOA classes ‘DNA Transcription’ and ‘DNA Replication’ were not significantly different anymore after 24h of treatment. Sigeti et al. found that for *Bacteroides fragilis*, metronidazole inhibited DNA synthesis after 20 min. It also inhibited RNA synthesis, but only after 60 min. Accordingly, secondary inhibition of DNA transcription occurs after inhibition of DNA replication (Sigeti, Guiney, and Davis 1983). Moreover, in *E. coli*, inhibition of either RNA synthesis and protein synthesis resulted in prevention of DNA replication (Maaløe and Hanawalt 1961). Furthermore, analysis of membrane integrity revealed an increase in cells with membrane damage with longer treatment time for most antimicrobials for *A. viscosus*. With *F. nucleatum,* this was only observed for some antimicrobials. However, the control showed a high relative abundance of cells stained with propidium iodide (i.e. damaged) after 3.5h of treatment (Supplementary 2). Propidium iodide has been demonstrated to be able to stain cells in the exponential growth phase (Shi et al. 2007) and *F. nucleatum* has been shown to reach the stationary phase only after 48h (Moreira Júnior et al. 2011). This could justify why increased membrane damage was only observed for a few antimicrobials with this strain.

Notably, polymyxin B was not grouping with its class (‘Membrane Disruption’) for *A. viscosus*, which can be explained by polymyxin B not being active against Gram-positive bacteria (Mohapatra, Dwibedy, and Padhy 2021). The phenomenon implies that an adapted pipeline to the one used in this experimental setup can be used for antimicrobial susceptibility testing. An antimicrobial with a known MOA that does not group with its respective MOA class implies that the compound is not active against the tested bacterium. This can be used in a fast assay where a range of compounds is screened to select the most effective antimicrobial. Moreover, it can be used to assess if a novel antimicrobial is effective against either Gram-positive or Gram-negative bacteria, or both.

A drawback of the presented workflow is that the difference in phenotypic fingerprints is mostly strain specific (Supplementary 5). Therefore, if novel compounds are assessed, it should be done on the same microorganism each time. To account for eventualities, the assessment should ideally be performed on multiple microorganisms. Notwithstanding, we found that grouping according to MOA could also be observed for a human saliva sample (Figure 3 (A)). The observation shows that the workflow can be robust for complex communities and suggests its potential for the personalization of antimicrobial treatment for patients. Hence, the selection of the most suitable treatment for the patient could be based on the response of the microbial community of the patient.

The performance of random forest classifiers was better for *F. nucleatum,* even though more statistically different comparisons between MOA classes were found for *A. viscosus*. As PhenoGMM looks for clusters of cells with similar properties inside the flow cytometry data, bins are set according to subpopulations found in the data (Ludwig et al. 2019; Rubbens et al. 2021b). We hypothesize that the resulting bins lead to distinctive features for each MOA class and that the random forest classification algorithm can select features that are unique for each MOA class. This is different from the statistical analysis, where only the similarity between MOA classes is compared to the similarity within each MOA class. Further investigation is needed to shed light on which factors lead to better classification performance and why differences are found between the used microorganisms.

When it came to predicting the MOA of an unseen compound, the shorter incubation time led to the best classification. Again, we argue that this may be the result of the phenotype of cells becoming more alike with increased treatment time because of cellular death. Remarkably, the classifier with the best performance (*F. nucleatum* after 24h of treatment) led to the worst prediction. Further investigation is needed, as this could be caused by multiple reasons. For example, the estimation of model performance could be biased, even though repeated k-fold cross validation is considered to be robust towards performance estimation (Jung 2018). Also, feature selection within the random forest model could be biased due to eventualities (*e.g.* a signal created by the interaction of the compounds with the medium) (Ribeiro, Singh, and Guestrin 2016). Alternatively, the (time-dependent) effect of the unseen compound simply resulted in a different phenotype of the bacterium as compared to the ones used for training.

The performance of the classifiers could be increased by expanding the database of phenotypic fingerprints used for training (Kwon and Sim 2013). Furthermore, the use of different classification algorithms, such as Kernel support vector machines, neural networks, or gradient-boosting decision trees, is worth investigating.

To our knowledge, only a few methods are able to reveal the MOA in a fast way. Bacterial cytological profiling is a fluorescence light microscopy-based technique that can be used to elucidate the MOA by pinpointing the molecular target of antimicrobials. The experimental procedure can be completed in a matter of hours. However, data analysis can be tedious as it involves image analysis (Lamsa et al. 2016; Nonejuie et al. 2013). Another fast technique is the use of AmpC reporter assays, but it can only be used to screen for inhibitors of cell wall sysnthesis (Sun et al. 2002). Our proposed workflow offers some clear advantages over other techniques. It is fast, cheap, and can be expanded to cover most MOA classes by including alternative staining methods. Finally, data analysis can be automated, leading to a reduced time to result.

To conclude, we found that the phenotypic fingerprints of bacteria treated with antimicrobials group according to the MOA of the antimicrobials. We found that this grouping could be observed both at the level of a single bacterial strain and at the level of a patient sample (*i.e.* saliva). Additionally, we showed that flow cytometry could be successfully used for the prediction of the MOA of novel antimicrobials and that a shorter treatment duration is favorable for this purpose. Thus, we provide proof of concept for a cheap and fast alternative for the elucidation of the MOA in the development of novel antimicrobials.

## Code and data availability

Raw flow cytometry data and metadata are available at FlowRepository under ID FR-FCM-Z6PB. The data analysis pipeline is available at https://github.com/famermans/FCM_MOA_Antimicrobials.

## Acknowledgements

The authors would like to thank Ruben Props, Frederiek-Maarten Kerckhof and Peter Rubbens for their advice during data analysis, Ioanna Chatzigiannidou for her advice during experimental design and Josefien Van Landuyt, Valérie Mattelin and Wannes Nauwynck for critically reading the manuscript. This work was supported by Research Foundation – Flanders (FWO, Belgium) (grant number: G0B2719N).

## Author contributions

FM and NB conceived and designed the study. FM and HB performed the experimental work. FM and HB analyzed the data. FM wrote the manuscript. CT, WT and NB supervised the findings of this work. All authors reviewed and approved the manuscript.

## Conflict of interest

The authors declare no conflict of interest.

